# Control-theoretic immune tradeoffs explain SARS-CoV-2 virulence and transmission variation

**DOI:** 10.1101/2021.04.25.441372

**Authors:** Anish A. Sarma, Aartik Sarma, Marie Csete, Peter P. Lee, John C. Doyle

**Affiliations:** Computation and Neural Systems, California Institute of Technology; Pasadena, CA 91105; Division of Pulmonology, Critical Care, Allergy, and Sleep Medicine, Department of Medicine, University of California-San Francisco; San Francisco, CA 94143; Department of Immuno-Oncology, City of Hope Comprehensive Cancer Center; Duarte, CA 91010; Control and Dynamical Systems, California Institute of Technology; Pasadena, CA 91105

## Abstract

Dramatic variation in SARS-CoV-2 virulence and transmission between hosts has driven the COVID-19 pandemic. The complexity and dynamics of the immune response present a challenge to understanding variation in SARS-CoV-2 infections. To address this challenge, we apply control theory, a framework used to study complex feedback systems, to establish rigorous mathematical bounds on immune responses. Two mechanisms of SARS-CoV-2 biology are sufficient to create extreme variation between hosts: (1) a sparsely expressed host receptor and (2) potent, but not unique, suppression of interferon. The resulting model unifies disparate and unexplained features of the SARS-CoV-2 pandemic, predicts features of future viruses that threaten to cause pandemics, and identifies potential interventions.

## Main Text

Variations in virulence and transmission, shorthanded as the dual puzzles of asymptomatic cases and superspreaders, have made SARS-CoV-2 infection and spread difficult to predict and control (*1*–*4*). The relationship between pathogen virulence and transmission has been a subject of longstanding speculation and formal study (*5*–*7*), and continues to be debated in the context of variation in SARS-CoV-2 infection (*3, 8*–*10*). The complexity of the immune response has impeded a unified mechanistic understanding of virulence, transmission, and variation, relevant to SARS-CoV-2 and future emerging viruses (**Figure 1**). Here, we extend techniques from control theory, a mathematical framework that has been used to analyze complex feedback systems in both engineered and biological settings (*11*–*14*), to immune biology to analyze SARS-CoV-2 virulence and transmission.

**Fig. 1.**
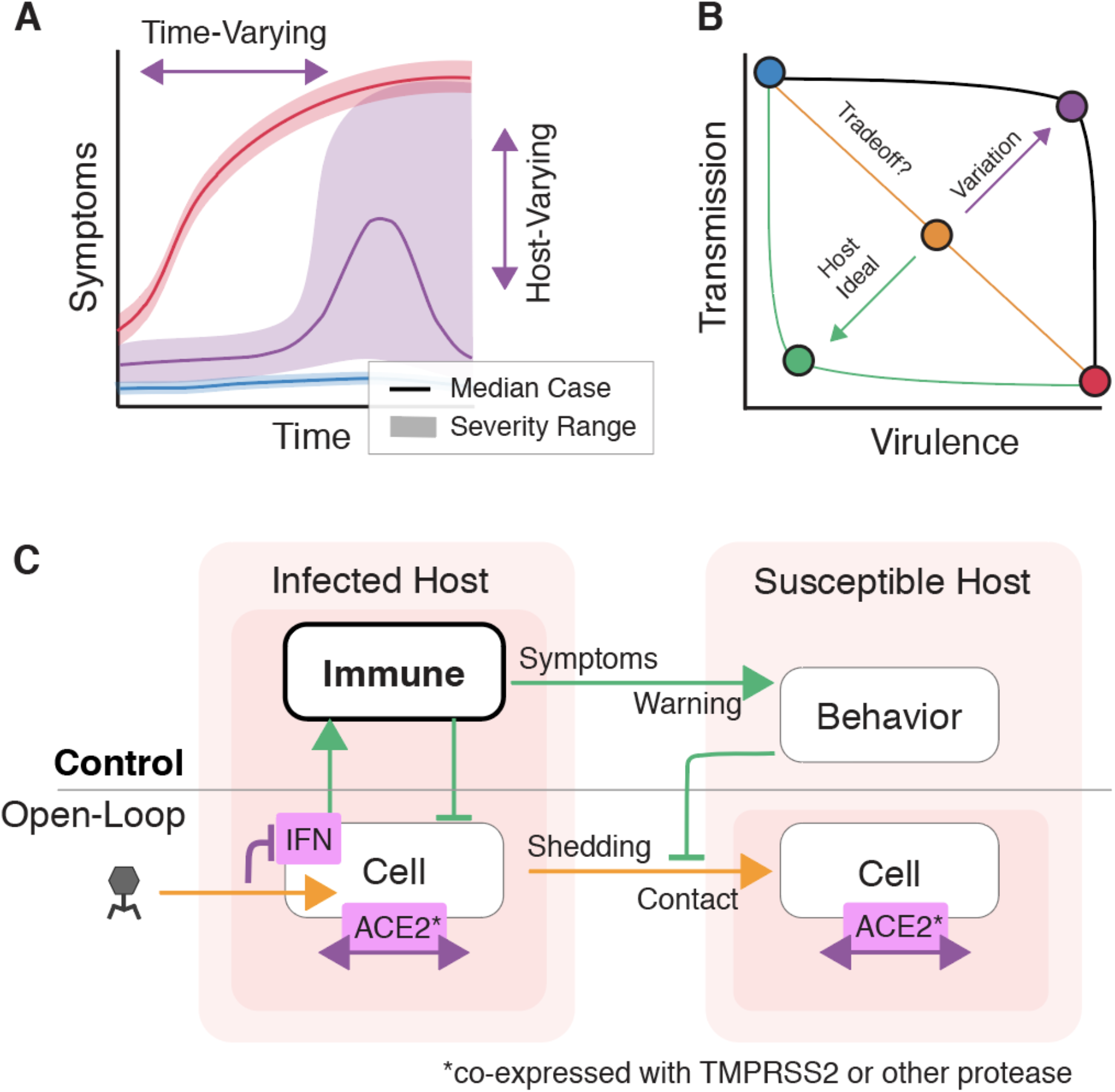
A control theory framework to analyze virulence and transmission. (A) Schematic time-series representations of symptoms for three viruses. The red and blue cases have relatively low variation across hosts and across time. The purple case, schematically like SARS-CoV-2, varies across hosts and across time, suggesting variation in host immune responses. (B) An apparent tradeoff between virulence and transmission can result from host immune responses. However, this tradeoff depends on host control mechanisms, and can be made more favorable to the host population or more favorable to the virus. (C) A block diagram shows the relationships between virulence and transmission control in two hosts, one infected with SARS-CoV-2 and one susceptible. The dynamics without control are shown in the lower half of the diagram, and the control responses in the upper half. Immune responses suppress shedding, create symptoms, and allow behavioral responses.

## Results

### A control-theoretic approach to immune dynamics

We use control theory to uncover mechanisms that lead to variation in virulence and transmission. Informally, we compute the best-case immune response, consolidating unmodeled immune dynamics into a control function *K* (**Figure 2A**). The best-case immune response minimizes virulence and implicitly suppresses transmission. We implement mechanistic details as constraints on the set of realizable control functions, and in this way identify mechanisms (constraints) for which even a best-case *K* yields virulence and transmission variation. This best-case *K* bounds any immune system model that we could have used, allowing us to pose rigorous questions without a detailed model of immune dynamics. Formally, we consider the robust control problem:

**Fig. 2.**
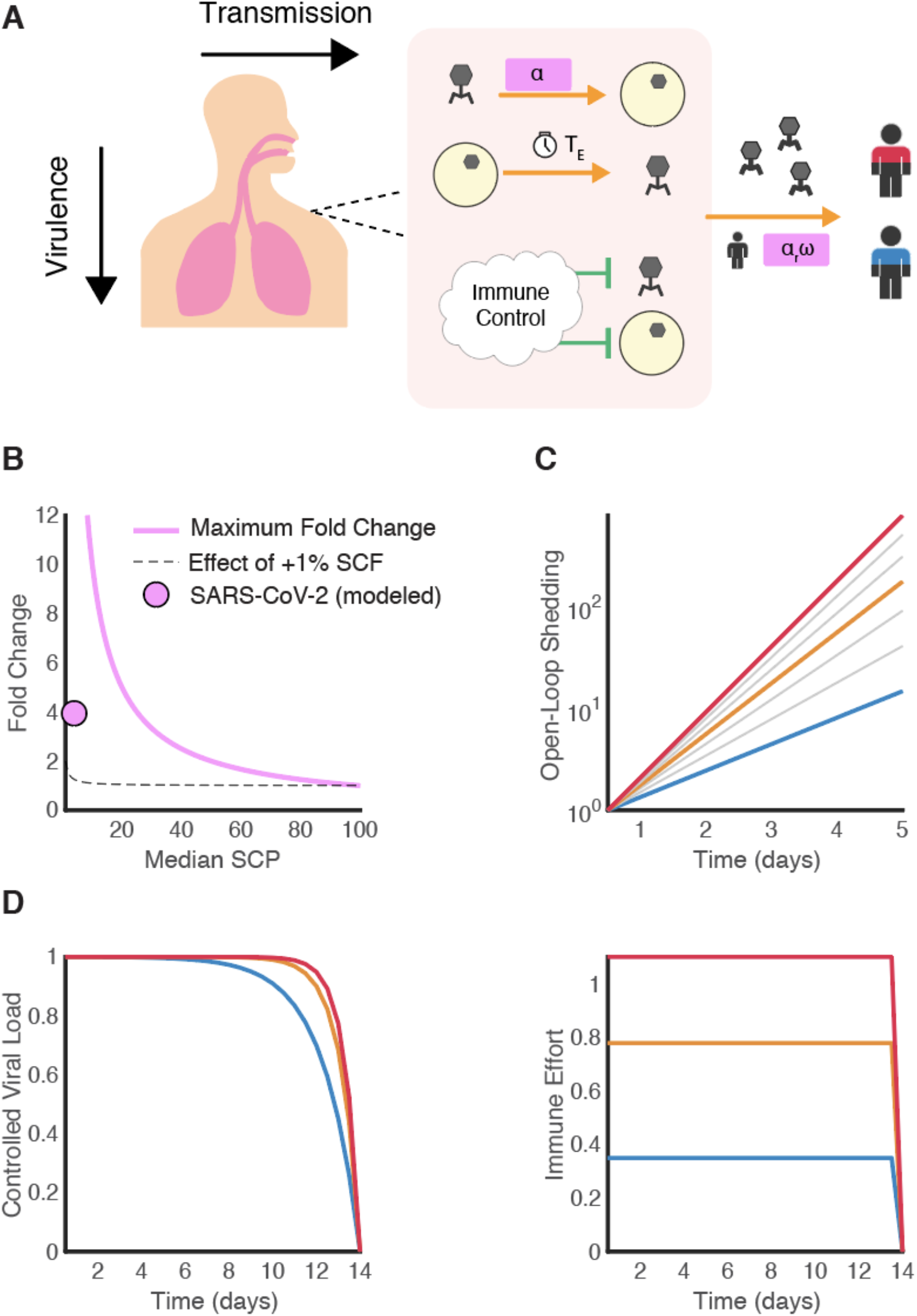
Host dynamics shape virulence and transmission. (A) Within each host, well-characterized kinetics govern viral replication. Viruses enter cells, replicate, exit after a delay, and degrade in the extracellular space. These kinetics are coupled to immune responses, for which we compute best-case bounds with control theory. (B) A low susceptible cell percentage (SCP) in the host enables variation. The maximum fold-change deviation from the median and the effect of a small fluctuation both grow as the median SCP approaches 0%. (C) An open-loop model removes all control elements and considers the underlying dynamics. Open-loop variation in viral shedding varies dramatically on relevant time-scales, amplifying variations in SCP. (D) Ideal extracellular immune control can create similar, low-variation viral load trajectories between hosts, but these similar trajectories require differing immune effort. The underlying open-loop dynamics directly shape virulence.

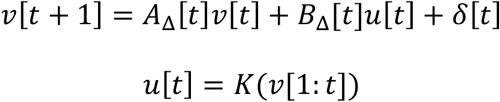

*v* is a vector of viral loads, *u* is the immune action, and new virus enters the system as *δ. A*_*Δ*_ and *B*_*Δ*_ are sets of time-varying matrices describing uncertain linearized dynamics. We leverage theorems guaranteeing that the best-case *K* always corresponds to a convex set, so that the best-case *K* computed from the set will be the best *K* over all realizable functions (*15*).

### The open-loop problem

We first consider the open-loop dynamics of viral replication, or equivalently *K* = 0. We model viral infection in the individual host as a three-step process: cell entry, replication in the cell, and release of virus from the cell after an eclipse period *T*_*e*_ (*16, 17*).

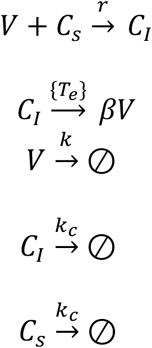

We derive *A* in terms of *α*, the number of productively infected cells that result from a single infected cell, where *(1-1/φ)* is the fraction of infected cells that constitutively turn over in a single eclipse period.

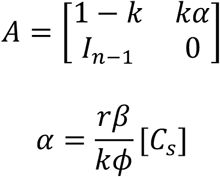

*α* scales linearly with [*C*_*s*_]. A small fraction of respiratory epithelial cells are susceptible to SARS-CoV-2 when compared to rhinovirus, respiratory syncytial virus, and influenza (*18*– *22*). A small susceptible cell fraction enables large *relative* variation consistent with reported single-cell data (*20, 21*) (**Figure 2B-C**).

### The closed-loop problem

We next consider the effects of immune control with innate extracellular effectors. Higher *α* requires stronger immune responses to achieve a comparable effect on viral load (**Figure 2D**). The immune response creates symptoms, which enable behavioral measures to avoid infection (*23*). We use a highly simplified model of avoidance and isolation, emphasizing the consequences of biological variation. We define transmission *R*_*CL*_, where *w(t)* is a warning signal and *γ(t)* = *exp(-pw(t))*. Initially, we take *w(t)* to be a scaled norm of the immune response, so that symptoms promote avoidance and isolation.

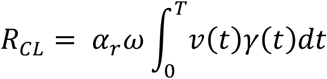

We extend this simple behavioral model to address a virus with a long presymptomatic period followed by uniformly severe infection (**Figure 3A-B**). Advance warning and isolation measures can contain such a virus. However, fully asymptomatic cases make advance warning more difficult. Fully asymptomatic cases need not be as contagious as presymptomatic-severe cases to have this effect, and low rates of fully asymptomatic cases can be tolerated (**Figure 3C**).

**Fig. 3.**
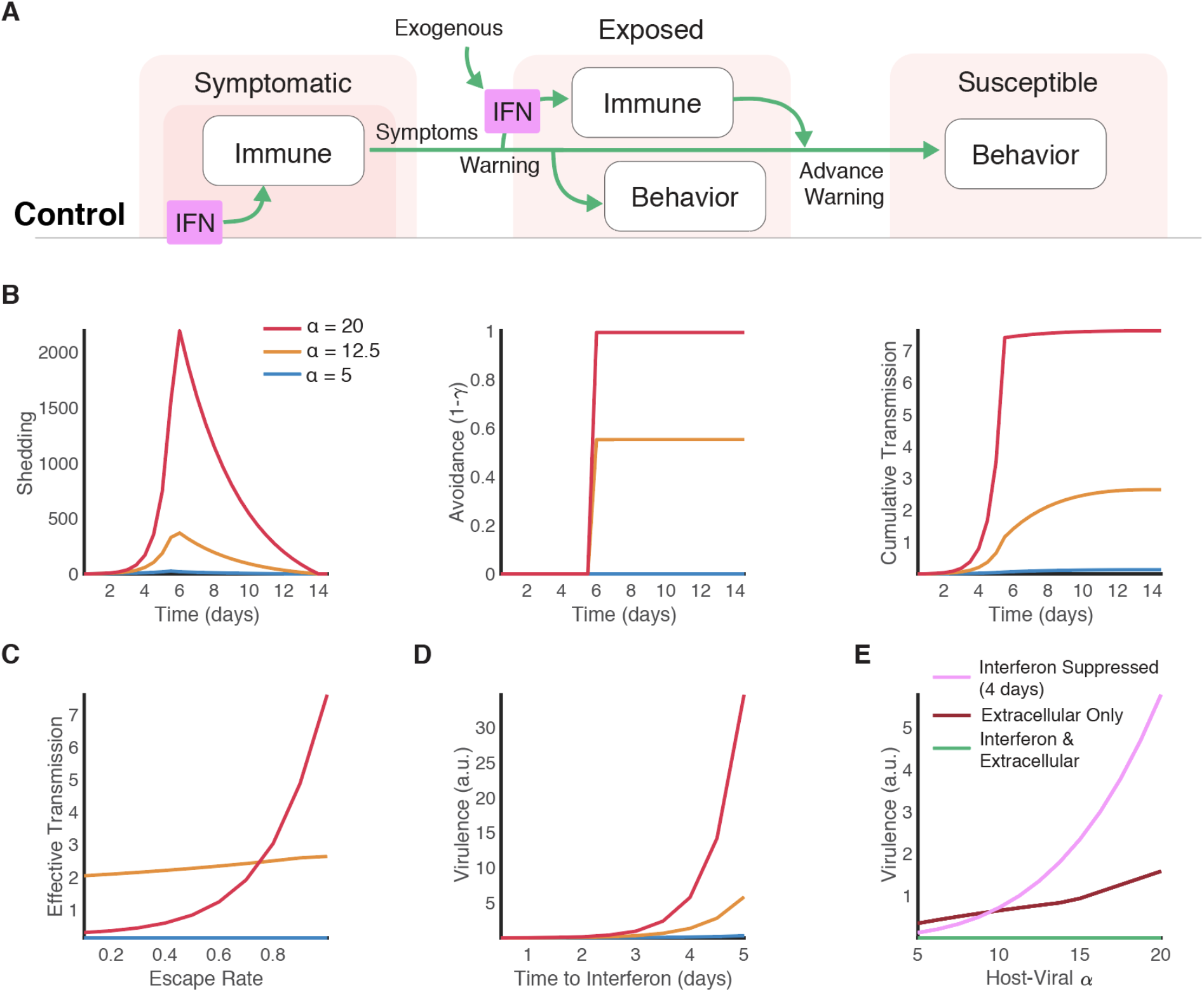
Layered control of virulence, transmission, and variation. (A) When all infected hosts are eventually symptomatic, advance warning allows isolation and potentially treatment measures in pre-symptomatic individuals. (B) Interferon-suppressed responses allow an extended period of viral replication and shedding during which avoidance behaviors are not possible (without advance warning). Transmission can be computed from the viral and immune trajectories. (C) Advance warning can reduce the effective transmission rate of presymptomatic individuals, but asymptomatic cases facilitate escape. As the rate of escape increases, the effective transmission from presymptomatic-severe individuals increases sharply. (D) Timing is a crucial determinant of interferon efficacy. Presymptomatic exogenous interferon administration can potentially reduce the eventual symptom burden in an individual who would otherwise experience severe disease. (E) Extracellular immune responses vary more with *α* than interferon-based immune responses, but interferon-suppressed responses vary most.

Interferon-based control varies less with *α* than extracellular responses, but interferon-suppressed control varies more. Early interventions with exogenous interferon can potentially reduce the eventual symptom burden in what would otherwise be severe cases (**Figure 3D-E**).

Taking these control layers together, we consider virulence and transmission as *α* varies. Presymptomatic-severe high-*α* cases take a dominant role in spreading the pathogen, especially where they interact with other high-*α* individuals (**Figure 4**).

**Fig. 4.**
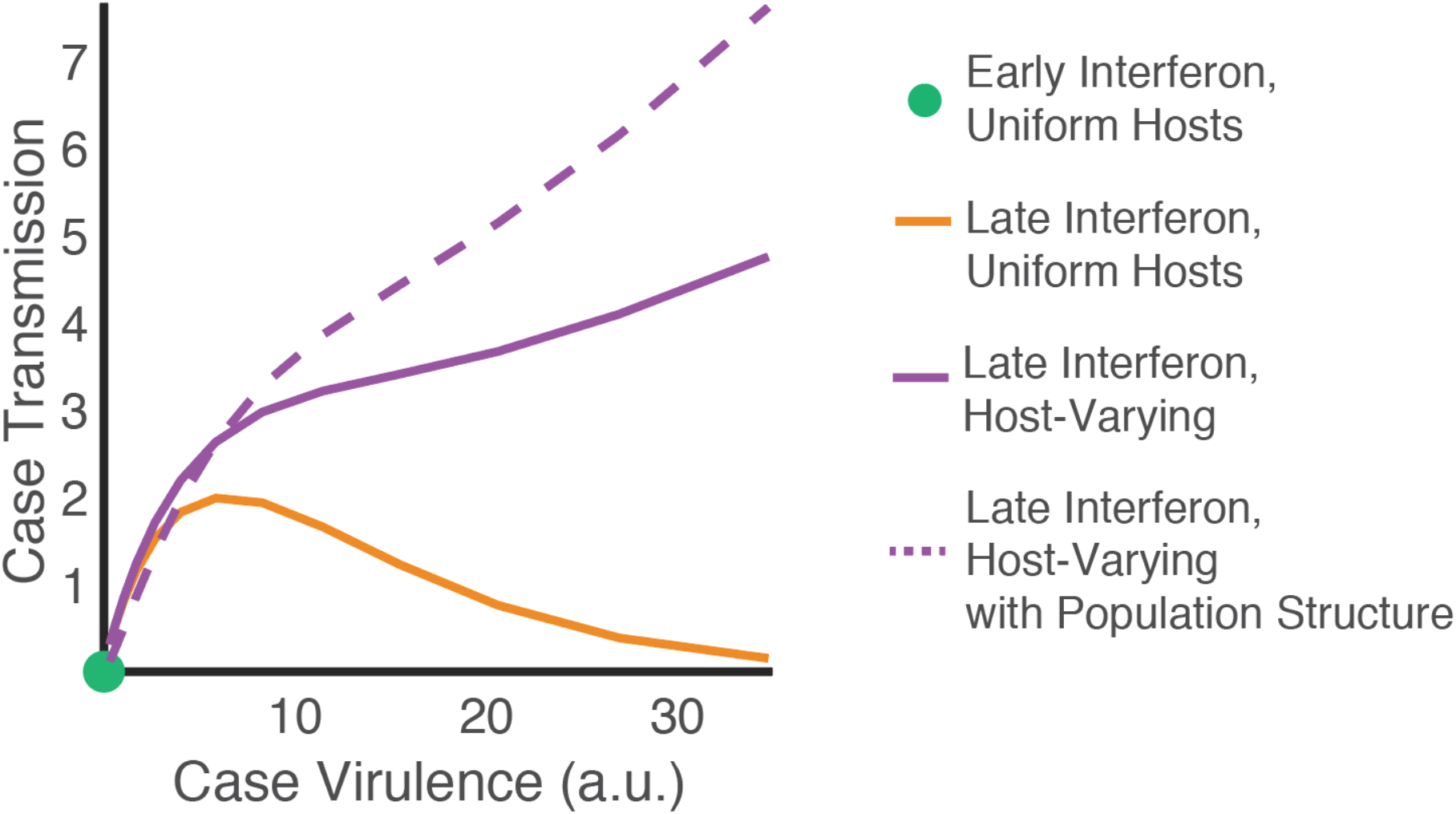
Virulence and transmission depend on host control strategies. The relationship between virulence and transmission depends on control conditions in individual hosts. If immune control is ideal for the host, replication is quickly blocked by interferon and neither serious symptoms nor substantial transmission occur. With interferon suppression, transmission peaks at low virulence. With interferon suppression and host variation, however, transmission is higher and peaks at higher virulence. This effect is amplified when high-*α* individuals interact, leading to both high presymptomatic shedding and high susceptibility.

## Discussion

### Other viruses

HCoV-NL63 and SARS-CoV-1 also bind to ACE2. HCoV-NL63 infection is asymptomatic or cold-like (*24*), while SARS-CoV-1 infection is typically severe, with some reported asymptomatic cases (*4, 25*). Viral infection in these cases could be biologically variable with median effects that are too mild or too severe to be evident in clinical outcomes. SARS-CoV-1 exhibits variable transmission, consistent with this interpretation (*26*). HCoV-NL63’s reduced virulence may result from a spike glycoprotein structure that decreases *α* (*27*).

### Interferon signaling

Clinical and experimental studies have shown that early exogenous interferon administration can reduce coronavirus infection severity (*28*–*32*). Our results suggest that presymptomatic interferon could be particularly beneficial in averting severe outcomes. Conversely, our results suggest a mechanism for harm from early immunosuppression in patients with SARS-CoV-2, consistent with clinical trial evidence (*33*). Because of the ubiquity of interferon suppression strategies in respiratory viruses, studies of control-guided interventions could facilitate responses to future emerging viruses.

## Notes

### Competing Interest Statement

The authors have declared no competing interest.

